# Predicting allergy and postpartum depression from an incomplete compositional microbiome

**DOI:** 10.1101/2025.02.28.640766

**Authors:** Andrey Shternshis, Bangzhuo Tong, Alkistis Skalkidou, Carolina Wählby, Dave Zachariah, Luisa W. Hugerth, Prashant Singh

## Abstract

Time series of compositional data are a common format for many high-throughput studies of biological molecules, e.g., analyzing the response to a treatment or with the aim of predicting an outcome. However, data from some time points may be missing, which reduces the size of the complete dataset. We propose a method for binary classification that includes imputation for missing values and logarithmic transformation of compositional data. Imputation approaches entail models that incorporate artificial data alongside true measurements, thereby supplementing the dataset. We consider two datasets from prospective analyses with associated target labels, aiming to improve prediction accuracy. We predict infants’ food allergies from their gut microbiome with a balanced accuracy of 0.72. We forecast postpartum depression based on gut microbiome data collected during pregnancy, with a balanced accuracy of 0.62. Features extracted from the microbiome time series, specifically ratios of bacterial abundance, are statistically significant indicators of depression.

## 1 Introduction

Compositional data are a common form of dataset in various fields, including geology (Thomas and Aitchison, 2005), demography (Bergeron-Boucher et al., 2017) and chemometrics (Korhoňová et al., 2009; Bosque-Sendra et al., 2012). In biology, essentially all data derived from sequencing are compositional, being limited to sequencing depth and generally deprived of absolute quantification (Quinn et al., 2018; Tsilimigras and Fodor, 2016). Compositional data lie within the probability simplex, meaning that all the elements are positive and sum to 1 (100%). This work focuses on classification problems where the input is a time series of compositional data and the target consists of binary labels (e.g., healthy, diseased). Time series of compositional data essentially arise when data from a study participant are collected periodically, for example, in gene expression profiling (Piening et al., 2018), exposomics (P. Zhang et al., 2021), and microbiome studies (Kindinger et al., 2017).

The goal of this study is twofold. First, we aim to develop an algorithm for effective binary classification when the input space consists of time points of compositional data. Second, we propose approaches for imputing missing values. Data may be missing at certain time points because of loss of follow-up, failure in sampling or DNA extraction. As a result, it is common in longitudinal studies to have a majority of participants with at least one time point with missing values. Therefore, techniques to impute artificial compositional data at missing time points to enable the use of incomplete data during classification are needed.

We propose imputation techniques and test them by applying them to two datasets. We specifically study microbiome data representing the fraction, or relative abundance, of various bacterial species. For the application part, the input space is chosen to be the gut microbiome, and the targets are postpartum depression for women and food allergies for children. Between 12% and 20% of mothers experience postpartum depression within the first three months after childbirth (Andersson et al., 2021; Gavin et al., 2005). (O’hara and Swain, 1996) identified a relationship between postpartum depression and low social support, stressful life events, and psychological disturbances during pregnancy. Identifying such predictors can help detect depression earlier and facilitate timely treatment (Fitelson et al., 2010). In addition to these predictors, we explore the potential of using the gut microbiome as a feature space for forecasting depression. Food protein-induced allergic proctocolitis (FPIAP) is a commonly recognized food allergy in early infancy that is diagnosed in 17% of cases (Martin, Virkud, Seay, et al., 2020). FPIAP is usually identified on the basis of a positive response to the removal of a food allergen. However, incorrect diagnoses can lead to unnecessary dietary changes for infants (Martin, Virkud, Dahan, et al., 2022).

The paper is organized as follows. First, we present an overview of methods applicable for classification of time points of compositional data. Features (e.g., species) important for classification may be located at different time points, as well as present changes in values over time. Second, we develop an approach for imputing missing compositional data, comparing various imputation methods. These methods are used to enlarge the input space for training classification models and to balance the dataset. For the datasets considered, imputation improves classification sensitivity and balanced accuracy. Finally, we demonstrated that the gut microbiome of pregnant women contains statistically significant features for classifying postpartum depression. For forecasting postpartum depression at six weeks after childbirth based on data collected during pregnancy, the balanced accuracy is 0.62, with a corresponding sensitivity of 0.39. For predicting FPIAP using data from the first six months of life, the sensitivity is 0.66, and the specificity is 0.77.

The next section reviews relevant papers on metrics, models, and applications. Section 3 introduces the datasets. Section 4 presents the methods for data transformation, imputation, and classification. Section 5 presents the results. Section 6 concludes the paper.

## 2 Related literature

(Brodersen et al., 2010) introduced balanced accuracy, which averages the accuracies for each target group. Given a confusion matrix for binary classification with the number of positives (*P*), true positives (*T*_*P*_), number of negatives (*N*), and true negatives (*T*_*N*_), the balanced accuracy is defined as follows:

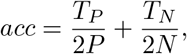

where *T*_*P*_ */P* is referred to as *sensitivity*, and *T*_*N*_ */N* is referred to as *specificity*. Random or constant decisions result in a balanced accuracy of *acc* = 0.5. In binary classification, balanced accuracy is equivalent to the area under the receiver operating characteristic curve (Bradley, 1997).

(Xia et al., 2013) and (Okazaki and Kawano, 2022) proposed a regression model for classification using logarithmic ratios of compositional data and a lasso regularizer for feature extraction (Hallac, Leskovec, and Boyd, 2015). Logarithmic ratios allow the transformation of compositional data into real space (Aitchison, 1982; Ibrahimi et al., 2023). Research on 24 datasets and a subset of compositional data transformations (Karwowska et al., 2025) concluded that transformations have a limited effect on classification accuracy.

(Martin, Virkud, Dahan, et al., 2022) analyzed the taxonomic differences in the gut microbiome between infants with food protein-induced allergic proctocolitis (FPIAP) and control cases (no FPIAP). The authors applied a random forest classifier to evaluate feature importance (Breiman, 2001) and determine prediction accuracy. When distinguishing control samples from three stages of FPIAP development (resolved, symptomatic, and presymptomatic), the balanced accuracy reached 0.53. In the case of the four groups, the balanced accuracy ranges from 0–1, with a value of 0.25 corresponding to a random guess. In a simpler scenario, when only samples from infants aged six months were used within the control or resolved groups, the balanced accuracy improved to 0.66, with a sensitivity of 0.76 and a specificity of 0.56.

(Andersson et al., 2021) used pregnancy- and childbirth-related variables along with psychometric questionnaires to predict postpartum depression. The authors achieved a balanced accuracy of 73% and a sensitivity of 72%. According to their findings, the most important variables for prediction include depression and anxiety during pregnancy as well as a history of depression. Postpartum depression primarily involves symptoms of anxiety and depression, which are closely correlated with changes in the gut microbiome (Y. Wang and Kasper, 2014). Moreover, several studies have reported changes in gut microbiome abundance in individuals with postpartum depression (Hao et al., 2020; S. Zhang, Lu, and G. Wang, 2023). (Tortajada et al., 2009) achieved a balanced accuracy of 83% and a sensitivity of 84% using background data, emotional alterations, and depressive symptoms. (Y. Zhang et al., 2021) obtained a balanced accuracy of 85% and a sensitivity of 82% using data extracted from electronic health records after childbirth.

## 3 Structure of the data and datasets

We present the analysis of the data in the following form: each participant’s data consists of *T* time points (tp) of compositional data. Measurements, represented as probability vectors (relative abundance summing up to 100%), are collected multiple times for each participant. Labels are associated either with individual time points or with the entire time series. Compositional data may be missing at certain time points. A sketch of the data is shown in Table 1. In the provided example, there are *n* data points, and we consider the case where *T* = 3. However, this is not a limitation for the classification and imputation approaches discussed in the next section.

**Table 1:**
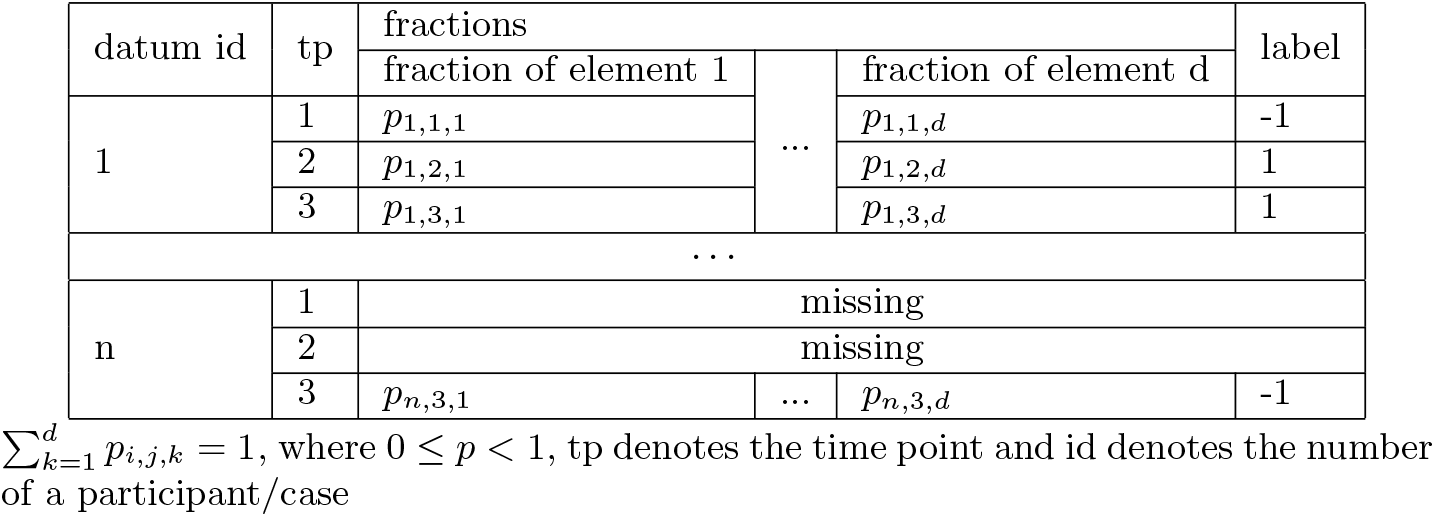
Sketch of the dataset.

The analysis is conducted on two datasets described below.

### Dataset 1

**The BASIC study** (Axfors et al., 2019) at Uppsala University Hospital collected data from approximately 5,000 pregnant women between 2009 and 2018. The participants were asked to complete the Edinburgh Postnatal Depression Scale (EPDS) questionnaire (Cox, Holden, and Sagovsky, 1987). Starting in 2016, gut microbiome data were included as part of the input space. Gut microbiome samples with fewer than 5 × 10^5^ reads were excluded from the analysis. Insufficient reads and missed appointments reduced the dataset size collected over the two years.

The compositional data represent relative species abundance, with a dimension of *d* = 713 species in the gut microbiome. Data were collected at three time points: during the 20th and 30th weeks of pregnancy and at six weeks postpartum. The outcome for each time point is the presence of depression, as defined by the EPDS. The label to forecast is at the last time point (*t* = *T*). For binarization, participants are considered healthy (*y* = −1) if their EPDS score is less than 12 and depressed (*y* = 1) otherwise (Wickberg and Hwang, 1996).

For complete data with no missing values, the dataset includes 82 data points with *y* = −1 and only 15 data points with *y* = 1, indicating a relatively small and imbalanced dataset. For data points where the last time point is missing, there are 16 positive and 108 negative labels. For a subset where the first or second time point is missing, there are 33 positive and 190 negative labels. A summary of the sizes is given in Table 2.

**Table 2:**
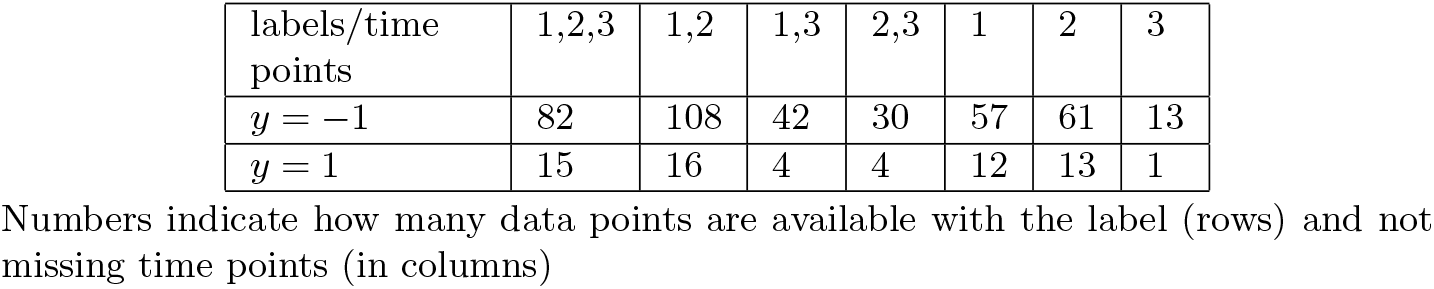
Amount of data points available for Dataset 1.

| labels/time points | 1,2,3 | 1,2 | 1,3 | 2,3 | 1 | 2 | 3 |
| --- | --- | --- | --- | --- | --- | --- | --- |
| $y = -1$ | 82 | 108 | 42 | 30 | 57 | 61 | 13 |
| $y = 1$ | 15 | 16 | 4 | 4 | 12 | 13 | 1 |

Numbers indicate how many data points are available with the label (rows) and not missing time points (in columns)

These data have been previously described in (Kimmel et al., 2025).

### Dataset 2: FPIAP

The second study examined the relationship between the gut microbiome and food allergies (Martin, Virkud, Dahan, et al., 2022). It involves 80 healthy infants (*y* = −1) and 82 infants diagnosed with food protein-induced allergic proctocolitis (FPIAP) during their first year of life (*y* = 1). The microbiome was sampled at multiple time points, specifically at the ages of 1, 2, 4, 6, 9, and 12 months. Table 3 summarizes the number of data points available at several time points. The amount of available data decreases as children approach one year of age.

**Table 3:**
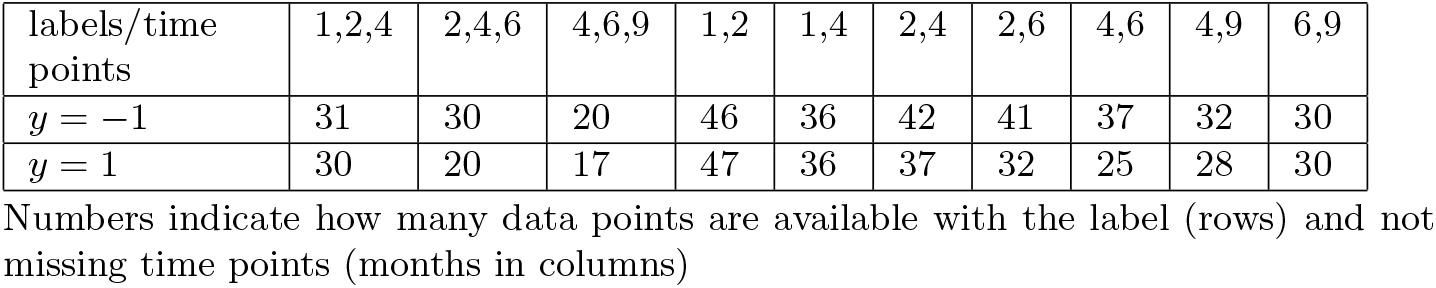
Amount of data points available for Dataset 2.

We omit three columns from the dataset that are associated with unassigned bacteria, reducing the dimensionality of the compositional data to *d* = 342.

## 4 Methods

Having discussed the format of the data, we now provide an overview of the methods applied for classifying data points by their target labels. This section begins with data preprocessing, which focuses on dimensionality reduction. Next, we discuss data transformations that enable us to work in real space instead of the original data format. We then introduce imputation techniques to handle incomplete time points, thereby enlarging the training set. Finally, we explore several classification approaches. When specific time points are used for classification, we assume that the data are either complete or that missing time points have been imputed via the methods outlined in this section.

### 4.1 Data prepossessing

The data are divided into testing, training, and validation sets with proportions of 0.2, 0.64, and 0.16, respectively. We use 5 validation and 5 testing sets for this partition. The imbalance ratio of labels remains consistent between groups across all sets.

Species with a sparsity greater than a predetermined threshold can be combined into a group called “other species.” Sparsity refers to the fraction of zero values across the training set calculated for each species. Depending on the threshold value, we can significantly reduce the dimensionality of the problem, although potentially important information may be lost by merging sparse species.

Finally, we identify the minimum value in the training set and replace all zero values with half of this minimum value, ensuring that the sum remains equal to 1. For the logarithmic transformations discussed later, all the elements must be positive. Alternatives for a data transformation that allows 0 values include (Scealy and Welsh, 2011; Firth and Sammut, 2023).

### 4.2 Data transformation

Currently, each data point is two-dimensional, containing compositional data at several time points. We consider four ways of transforming the data:

- *Compositional*. Each element *x*_*i*,*t*_ is considered an input feature, where 1 ≤ *i* ≤ *d* represents the number of species and 1 ≤ *t* ≤ *T* represents a time point.
- *Centered* log ratios (CLR, (Egozcue et al., 2003)). Each element at a time point is divided by its geometric mean. The logarithm of these ratios is called CLR: 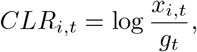 where 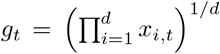. This transformation shifts the data from a compositional structure to a real space.
- *All* log ratios (ALR). We flatten data points to compositional data by concatenating time points and dividing by *T* . 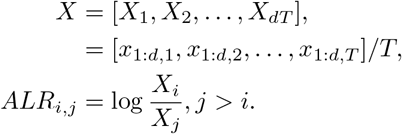 In this way, we increase the feature space dimension from *D* = *dT* to 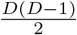 . By suggesting this transformation, we assume that the labels may not be determined by fractions at each time point but rather by changes in the fractions over time. A dummy constant element may be introduced to add logarithms of compositional elements to the input space together with all ratios.
- *Pivot* log ratios (PLR). The log ratios between clusters of elements are used here. The number of all possible clusters is large; thus, we limit it by pivot features, as suggested in (Fišerová and Hron, 2011). The clustering metric is the minimum variance within a cluster. It is known as the Ward method (Ward Jr, 1963). The PLR is the ratio of the geometric means of the left and right subtrees for each node. We set the first feature to be the geometric mean, *g*(*X*). As shown in Figure 1, the second feature is the ratio of the geometric means of the left (green) and right (orange) trees. All other subtrees are defined iteratively. With the PLR transformation, we do not increase dimensionality and retain the same amount of information as in the original dataset. Furthermore, it is possible to identify the specific clusters of bacteria used to calculate each feature.

**Figure 1.**
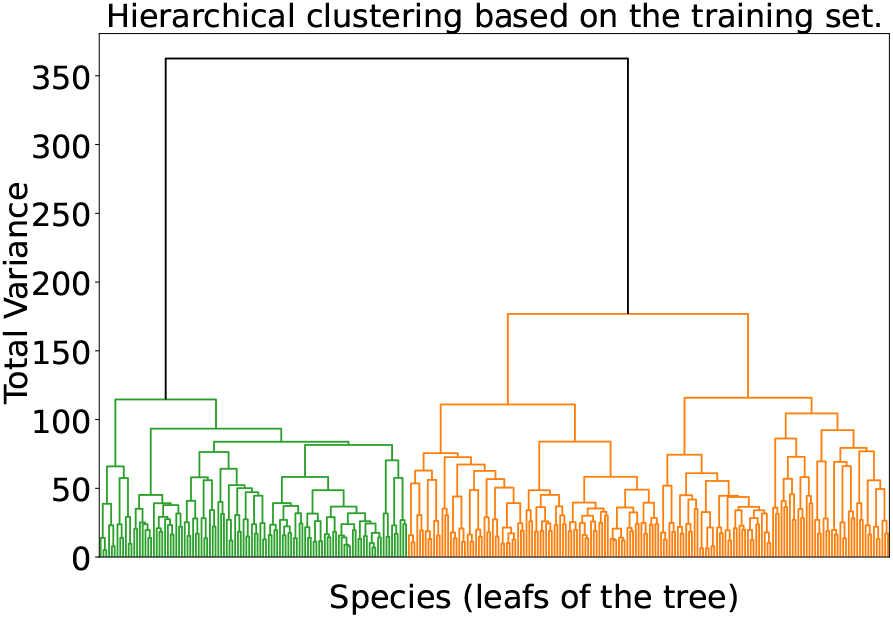
Hierarchical tree generated via the Ward method. Species merging was used.

### 4.3 Imputation

Imputation refers to the process of replacing missing data with artificially generated values. We employ five imputation approaches that use available time points to restore the missing points. A common step for all approaches is to transform the compositional data at each time point into a CLR. This transformation has a well-known inverse, called Softmax (Aitchison, 1982). The approaches differ in how they select the best-fit simulated predictions for imputation. In other words, we present various models with different objective functions for optimization.

- A matrix *A* and a vector *b* are chosen to minimize the least squares error between *AC*_*n*_ + *b* and *C*_*m*_, where the output *C*_*m*_ represents a missing time point and the input *C*_*n*_ represents time points that are not missing. The training set is used to find the optimal parameters. We consider linear regression as a baseline for comparison with more sophisticated models. A simpler model could be to substitute equal values instead of all missing fractions.
- Support Vector Regression (SVR, (Drucker et al., 1996)). SVR is a non-linear regression technique that aims to find a function that best fits the data by minimizing the prediction error within a specified margin. The optimization does not depend on the input dimensionality, allowing SVR to handle complex nonlinear relationships in the data.
- Gaussian process regression (GPR, (Williams and Rasmussen, 1995)). The approach provides a probabilistic approach that considers data as a realization of a stochastic process. The values imputed have the maximum likelihood. GPR treats models as distributions over functions. Predictions are made via the mean of the Gaussian process.
- Conditional Variational Autoencoder (CVAE, (Sohn, Lee, and Yan, 2015)). Another probabilistic approach allows the restoration of missing values from white noise. During training, noise is encoded from all given time points. The prediction is decoded from inputs and a noise term.
- Conditional generative adversarial network (cGAN, (Mirza and Osindero, 2014)). Another way to learn a generative neural network. In this paradigm, the goal is to create predictions indistinguishable from real data by a discriminator. Similar to the previous approach, predictions are generated from a noise term and available data allowing for nonuniqueness of the imputed values.

See Figures 2 and 3 for the architectures of Neural Networks. To limit the number of parameters to train due to the small dataset size, we consider neural networks with one hidden layer for the last two models. See section 4.5 for further implementation details.

**Figure 2.**
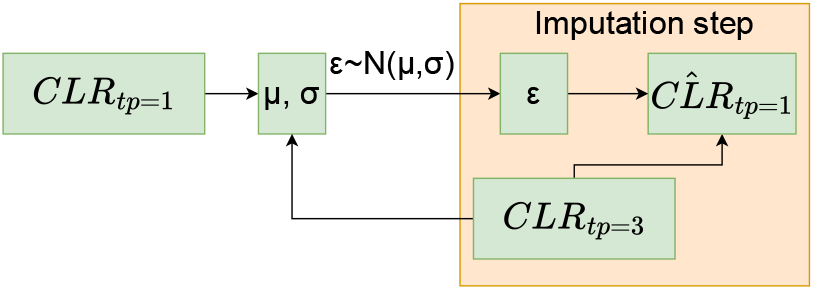
Example of the CVAE architecture where tp 1 is reconstructed from tp 3 and random noise *ϵ*. During imputation, *ϵ* is sampled from Gaussian *N* (0, 1).

**Figure 3.**
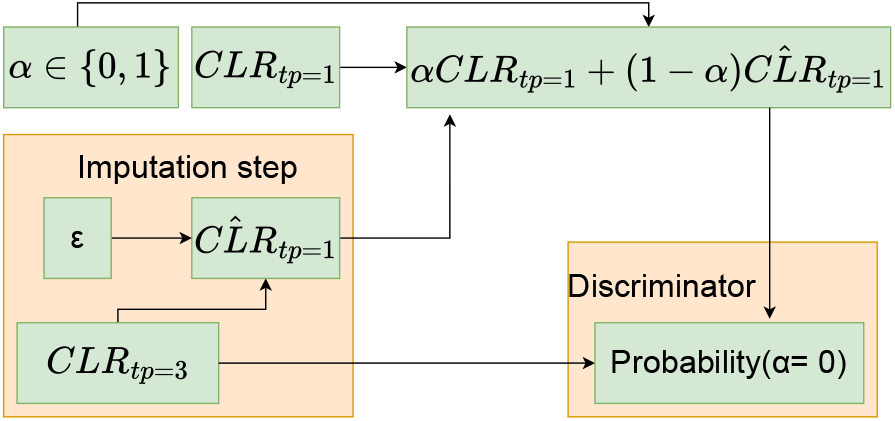
Example of cGAN architecture where tp 1 is reconstructed from tp 3 and random noise *ϵ, ϵ* is sampled from Gaussian *N* (0, 1).

### 4.4 Classification

We select random forest (Breiman, 2001) as a commonly used classifier (Boulesteix et al., 2012; Qi, 2012). We compare it with the support vector classifier (SVC, (Cortes and Vapnik, 1995; Noble, 2006)) and adaptive boosting (AdaBoost, (Freund and Schapire, 1997)), which are known state-of-art methods for binary classification. Both random forest and AdaBoost are based on decision trees. While random forest aggregates the classifications made by individual decision trees, AdaBoost adjusts sample weights based on the classification outcomes of the decision trees. Approaches that explicitly control classification errors conditioned on a class are also considered, as discussed in (Care, Ramponi, and Campi, 2018). The classification is performed on a subset of the feature space, with the selected features determined by analysis of variance (ANOVA) (St, Wold, et al., 1989).

### 4.5 Implementation and hyperparameters

Scripts to reproduce this research are available at the following GitHub link: github.com/AndreyShternshis/prediction-and-imputation-for-microbiome. The remainder of this section is dedicated to the hyperparameters chosen for the experiments.

- We reserve 20% of the data for the test set. The remaining data are divided into 5 cross-validation sets. This results in 3 and 2 data points with positive labels for the test and validation sets in Dataset 1, respectively.
- The threshold for sparsity is 0.5 for Dataset 1 and 0.9 for Dataset 2. That is, we combine bacteria into one group for dimensionality reduction if their sparsity is greater than one half for the dataset regarding depression.
- To reduce the execution time, we limit the number of features to 20, as the optimal number of features rarely exceeds 10 for the selected methods.
- During the imputation step, the size of the latent space for neural networks, which is used to generate random noise, is set to 1. We use a batch size of 1, with a maximum of 1000 epochs. The stopping criterion is based on an increase in validation error, meaning that the “patience” parameter is set to 0. To train the neural networks, the default settings of the Adam optimizer (Kingma and Ba, 2014) from the torch.optim library (Paszke et al., 2017) are used.
- For the lasso regularizer, we set a random state of 10, an *L*_1_-norm weight of 10, and increase the maximum number of iterations to 10^4^ to ensure convergence of the total loss.

## 5 Results

We start by testing log-transformation on a complete part of the dataset regarding postpartum depression. Log transformations are used for converting input compositional data from the (0, 1) range into the real number space. We then proceed with forecasting the depression by analyzing a reduced feature space and applying imputation approaches in section 5.2. The results of log-transformation and imputation techniques for the allergy dataset are given in section 5.3.

### 5.1 Classification of depression

First, we present the results of a classification problem using data points from Dataset 1 that have no missing time points, including the last one after childbirth, which is available for analysis. We apply data transformation through log ratios, as described in section 4.2. We also examine whether dimensionality reduction by merging bacteria, as detailed in section 4.1, improves performance.

Given that the data are imbalanced, we investigate whether assigning different weights during classification enhances accuracy. The optimal number of features, *n*, is selected through cross-validation. The balanced accuracies from the validation sets are averaged using either the mean or median. The median is more robust to random perturbations, often resulting in a smaller optimal number. The results are averaged over 5 runs of the classifier with different random seeds. The mean results from the 5 testing sets, including balanced accuracy (acc), sensitivity (sens., correctly identified depression), and specificity (spec., correctly identified no depression), are presented in Table 4. The p-values in the tables test the hypothesis that the balanced accuracies on the testing sets are greater than 0.5. Thus, a one-tailed test for the mean accuracy is used.

**Table 4:**
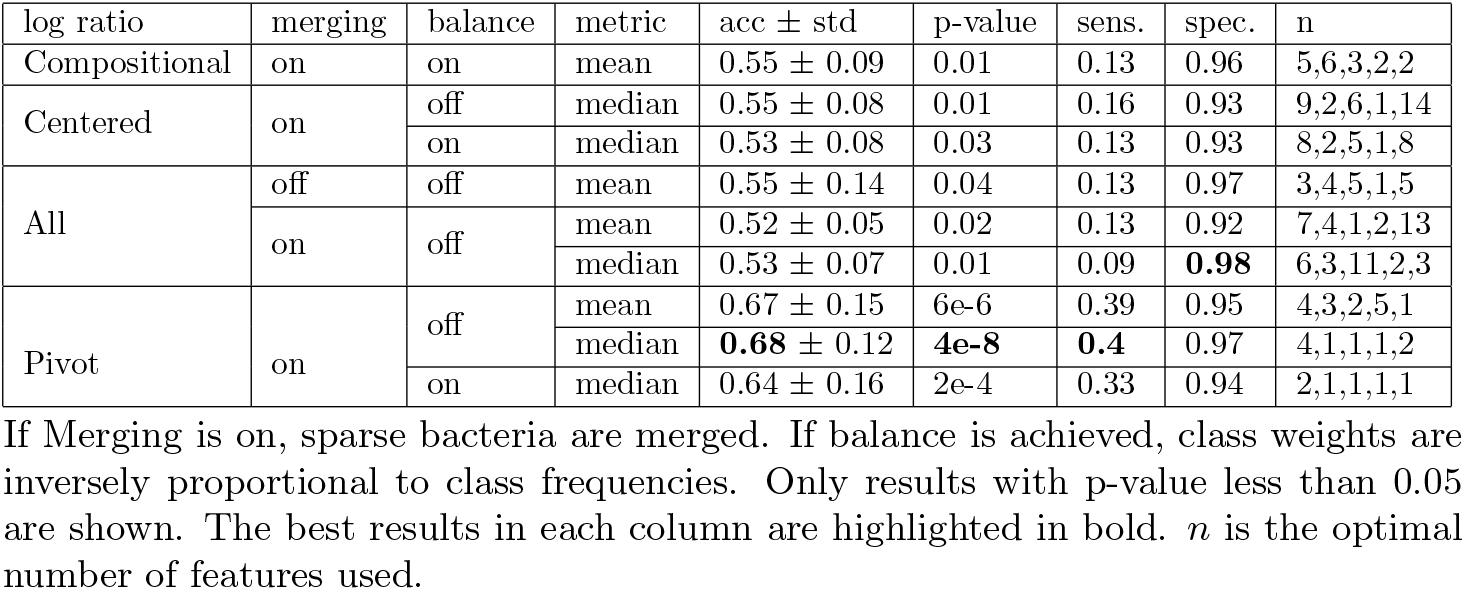
Accuracy for different types of log ratios for Dataset 1.

The pivot log ratio, which considers differences between clusters of species, outperforms other transformations under analysis. Pivot log ratios allow us to look at the data in terms of differences in bacterial abundance as well as changes over time while maintaining the original dimensionality. We conclude that the highest accuracy is achieved with dimensionality reduction by merging species. Adjusting weights according to class size does not appear to be beneficial. These choices for merging (on) and balancing (off) are fixed for the remainder of the study. Other user options, such as classifier selection and feature selection, are detailed in Appendix A.

### 5.2 Forecasting depression

Four out of the five test sets show a balanced accuracy, averaged over five runs of classification, greater than 0.5. For these four runs, we present the features used to classify depression. It appears that all features are ratios between two bacteria. The relationship between these features and the labels is clearly identified in the case of pivot log ratios but not with all log ratios because of the dimensionality of the feature space introduced by the log transformations. We present a feature list when the metric is chosen as the mean, which provides a more comprehensive representation. Note that the best results, in terms of balanced accuracy and sensitivity, were obtained using the median to determine the optimal number of features. In this case, four features are selected, as specified in Table 5.

**Table 5:**
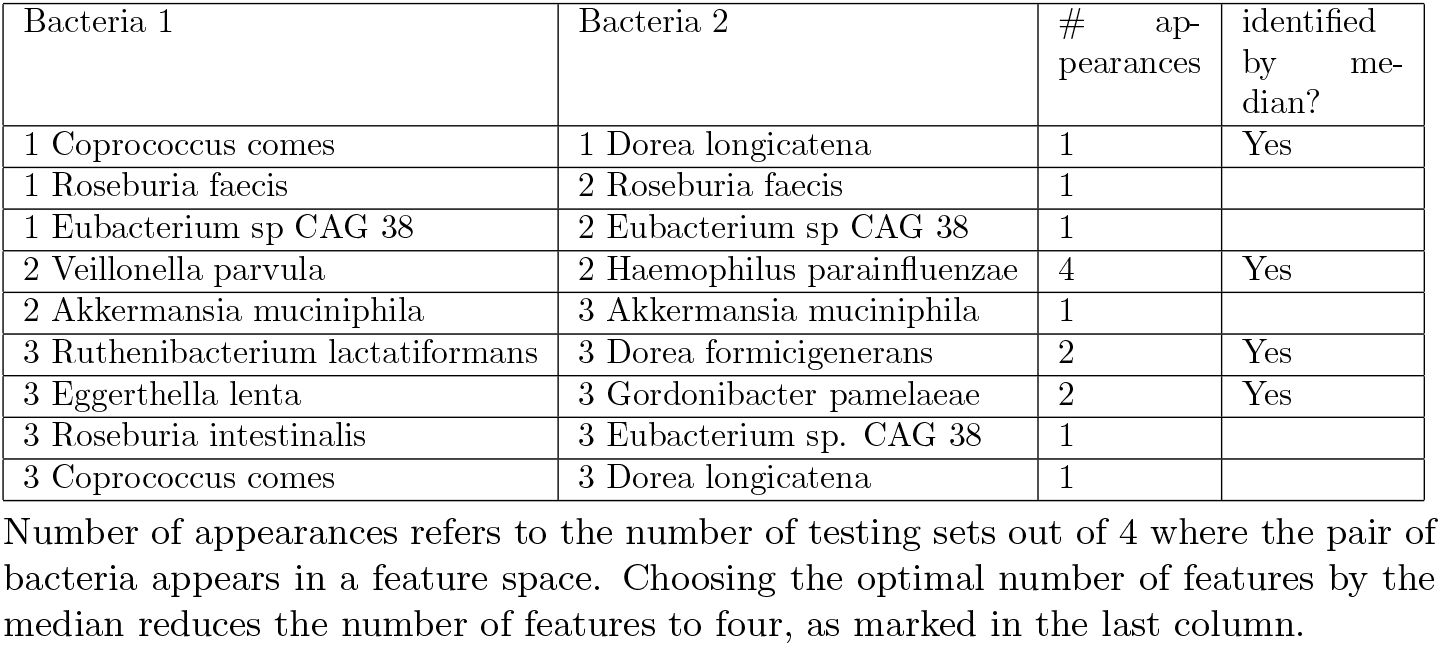
List of bacteria used in the classification of depression.

Note that only half of the bacterial species from Table 5 are detected during the last time point (postpartum). Across all four test sets, the same important feature for classifying depression is the difference between the abundances of the two bacteria Veillonella parvula and Haemophilus parainfluenzae at time point 2. Some features represent changes in the abundance of the same bacteria over time. The same pair of bacteria also appears to be important at both time points 1 and 3. Below are some further results based on these features:

- Using only the four selected features, we obtain a sensitivity of 0.33 with a specificity of 0.993. When only the first two features, which can be obtained during pregnancy, are used, the specificity improves to 0.995.
- However, when we included the analysis data points with missing data at time point 3 (women who missed the last visit), the balanced accuracy decreased to 0.55, with a sensitivity of 0.17. The result is still statistically significant, with a p value of 4*e*− 4. That is, our forecasting works better under the assumption that a woman under consideration does not miss the last visit after a childbirth.
- Taking the ratio between the abundances of Veillonella parvula and Haemophilus parainfluenzae at two time points during pregnancy as two features, we increased the balanced accuracy to 0.61, with a sensitivity of 0.25. Previous studies have shown that Veillonella parvula is positively correlated with major depressive disorder (X. Zhang et al., 2022) and bipolar disorder (T. Huang et al., 2023), whereas low levels of Haemophilus parainfluenzae were associated with psychotherapeutic responses in (Malan-Müller et al., 2024).
- When we augment the training dataset with imputed values from Gaussian process regression (GPR), we do not observe an improvement in classification accuracy. The results for various imputation methods are summarized in Table 6. These results are obtained from five imputation sets for each testing set, as the results are random due to parameter initialization and random noise generation. Multiple rounds of imputation highlight the uncertainty around the best-fitting imputed values (Rubin, 2004). We acknowledge that incomplete datasets with imputed values may have distributions that differ from those of complete datasets. To address this, we augment the input space with binary class variables indicating whether the data at each time point were imputed. These variables do not affect the prediction results.
- However, applying imputation with the goal of balancing the dataset increases accuracy metrics. When incomplete time points with only positive labels are included, the balanced accuracy increases to 0.62, with a sensitivity of 0.39.

**Table 6:** Accuracy for different types of imputation for Dataset 1.

| imputation | tp imputed | acc ± std | p-value | sens. | spec. | mean n |
| --- | --- | --- | --- | --- | --- | --- |
| linear regression | 1 | $0.56 \pm 0.06$ | 7e-05 | 0.15 | **0.97** | 1.8 |
| SVR | 2 | $0.53 \pm 0.06$ | 3e-3 | 0.16 | 0.91 | 1.6 |
| | 1 | $0.58 \pm 0.06$ | 1e-06 | 0.19 | 0.96 | 1.8 |
| | 1,2 | $0.55 \pm 0.06$ | 4e-4 | 0.2 | 0.9 | 1.6 |
| GPR | 2 | **$0.61 \pm 0.1$** | 3e-23 | **0.27** | 0.95 | 2 |
| | 1 | $0.6 \pm 0.04$ | 1e-55 | 0.23 | 0.96 | 1.8 |
| | 1,2 | $0.58 \pm 0.08$ | 6e-22 | 0.25 | 0.91 | 1.6 |
| CVAE | 2 | $0.59 \pm 0.11$ | 8e-16 | 0.23 | 0.94 | 1.88 |
| | 1 | $0.57 \pm 0.07$ | 4e-20 | 0.19 | 0.96 | 1.8 |
| | 1,2 | $0.57 \pm 0.08$ | 1e-18 | 0.2 | 0.95 | 1.76 |
| cGAN | 2 | $0.6 \pm 0.09$ | 2e-23 | 0.25 | 0.96 | 2 |
| | 1 | $0.59 \pm 0.07$ | 2e-27 | 0.21 | 0.96 | 1.8 |
| | 1,2 | $0.58 \pm 0.08$ | 3e-20 | 0.22 | 0.94 | 1.72 |

Imputing tp 1 and tp 2 from each other is considered. Imputing tp 2 by linear regression gives 0 values of bacterial abundance. The best results in each column are highlighted in bold. $n$ is the optimal number of features used.

To balance the dataset, we select the GPR for imputing missing time points, as it preservers the same accuracy according to Table 6. This can be interpreted as follows. GPR produces the most realistic time points, according to its estimated likelihood, since testing sets, where the accuracy is calculated, contain only complete time points.

We also found that missing data was not a significant predictor of depression. By using two binary variables denoting the presence of missing values at the time points during pregnancy, we achieve a balanced accuracy of 0.5. In contrast, using depression states during pregnancy results in a balanced accuracy of 0.59, with a sensitivity of 0.24. We outperform this result by using the gut microbiome.

### 5.3 Prediction of food allergies

By presenting the second dataset, we aim to demonstrate the effectiveness of imputation approaches for prediction tasks and provide a more comprehensive comparison of the different methods. We fix the number of time points and consider three consecutive time points: 2, 4, and 6 months. By merging bacteria at these time points, we achieve a balanced accuracy of 0.58. Applying a logratio transformation to the input space increases the balanced accuracy to 0.66, with a sensitivity of 0.52. When the training set is augmented by data points with one-third of the time points imputed by the cGAN, the balanced accuracy improves to 0.72, with a sensitivity of 0.66. For a detailed comparison of the transformation and imputation approaches, see Tables 7 and 8, respectively.

**Table 7:** Accuracy for different types of log ratios for Dataset 2.

| log ratio | metric | acc ± std | p-value | sens. | spec. | n |
| --- | --- | --- | --- | --- | --- | --- |
| compositional | mean | $0.59 \pm 0.07$ | 1e-06 | 0.36 | **0.81** | 2 14 1 18 14 |
| | median | $0.64 \pm 0.08$ | 2e-8 | 0.5 | 0.77 | 1 1 1 9 2 |
| All | mean | **$0.66 \pm 0.07$** | **2e-11** | **0.52** | 0.79 | 16,13,6,14,7 |
| | median | $0.59 \pm 0.12$ | 4e-4 | 0.39 | 0.79 | 5 6 6 6 2 |
| CLR | mean | $0.55 \pm 0.1$ | 0.01 | 0.32 | 0.77 | 2 18 6 20 2 |
| | median | $0.59 \pm 0.12$ | 3e-4 | 0.44 | 0.75 | 7 10 4 10 2 |
| PLR | mean | $0.57 \pm 0.18$ | 0.04 | 0.49 | 0.65 | 6 1 13 9 14 |

Only results with p-value less than 0.05 are shown. The best results in each column are highlighted in bold. $n$ is the optimal number of features used.

**Table 8:** Accuracy for different types of imputation for Dataset 2.

| imputation | data points imputed | acc ± std | p-value | sens. | spec. | mean n |
| --- | --- | --- | --- | --- | --- | --- |
| linear regression | all | $0.59 \pm 0.15$ | 5e-3 | 0.57 | 0.61 | 14 |
| | with 1 tp missing | $0.59 \pm 0.07$ | 2e-6 | 0.59 | 0.58 | 7.4 |
| SVR | all | $0.61 \pm 0.12$ | 5e-5 | 0.59 | 0.63 | 8 |
| | with 1 tp missing | $0.56 \pm 0.12$ | 0.01 | 0.52 | 0.59 | 6.8 |
| GPR | all | $0.64 \pm 0.13$ | 7e-23 | 0.55 | 0.73 | 8.8 |
| | with 1 tp missing | $0.67 \pm 0.08$ | 4e-49 | 0.65 | 0.68 | 8 |
| CVAE | all | $0.56 \pm 0.13$ | 1e-06 | 0.53 | 0.59 | 8.92 |
| | with 1 tp missing | $0.65 \pm 0.12$ | 4e-28 | 0.63 | 0.68 | 5.68 |
| cGAN | all | $0.66 \pm 0.12$ | 5e-29 | 0.58 | 0.75 | 8.6 |
|  | with 1 tp missing | **$0.72 \pm 0.11$** | 3e-43 | **0.66** | **0.77** | 6.96 |

The best results in each column are highlighted in bold. For "all" data points, one or two time points can be missing. In "with 1 tp missing", two other time points are available. $n$ is the optimal number of features used.

According to the study by (Martin, Virkud, Dahan, et al., 2022), compositional data are informative with respect to food allergies. In our analysis, we present results for imputation techniques applied to compositional data without log-transformation. By increasing the size of the training set through the inclusion of incomplete data points, we improve the prediction accuracy. We identified 20 bacteria used for allergy prediction during testing, half of which appeared multiple times across the 5 testing sets and 5 seeds for imputation. Notably, the abundance of Lactobacillus is an important feature for classification at all three time points. Additionally, measures of the abundance of the Lachnospiraceae family at time points 1 and 3 are also key predictors. Both features were identified in the research (Martin, Virkud, Dahan, et al., 2022).

To validate the choice of time points, we repeat the imputation and prediction for two alternative sets of months where the microbiome samples are taken: [1, 2, 4] and [4, 6, 9]. The results shown in Figure 4 demonstrate that the highest balanced accuracy and sensitivity are achieved with the time points at 2, 4, and 6 months. This outcome can be explained by a stronger correlation between the abundance of certain bacteria and food allergies, as well as the lack of complete data for months [4, 6, 9] for training.

**Figure 4.**
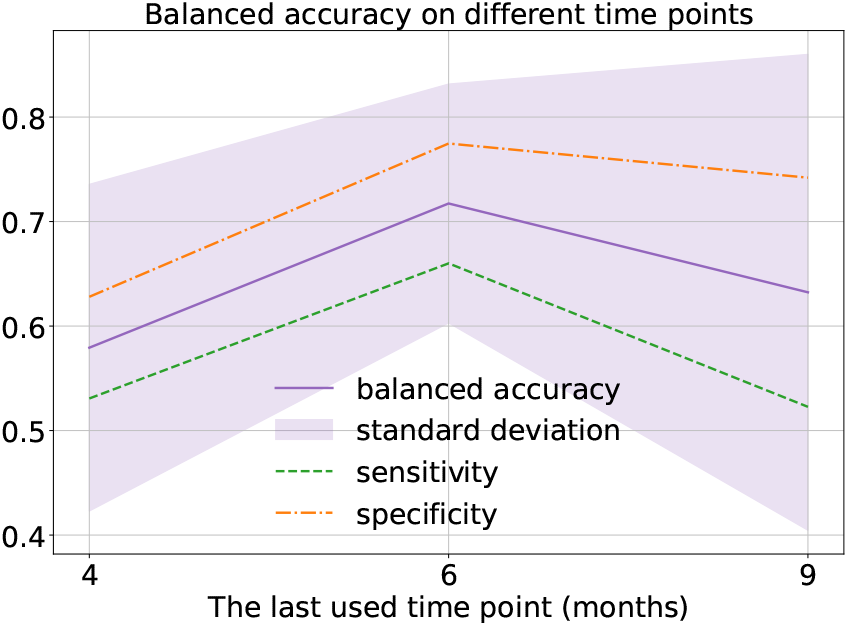
The prediction results for Dataset 2. The x-axis represents the last time point considered in months of life. The y-axis represents metrics such as balanced accuracy, sensitivity and specificity.

## 6 Discussion

We have presented an approach for binary classification when several time points of compositional data are given. The implementation of the approach is publicly available for reproducibility and further use. We construct the following algorithm for the prediction task when the input is microbiome data. We (i) reduce the dimensionality and sparsity of the data, (ii) impute missing data and merge it with the training set, (iii) transform the data via log ratios, (iv) select the features via analysis of variance on a training set (St, Wold, et al., 1989), and (v) use the features as the inputs of a random forest classifier (Breiman, 2001). The optimal number of features is chosen via cross-validation. The results are averaged on the testing sets, and imputation inference and initialization parameters are used.

We have empirically shown that the Lasso regularizer (Hallac, Leskovec, and Boyd, 2015) used in (Xia et al., 2013; Okazaki and Kawano, 2022) yields classification results that are worse than those of ANOVA for our choice of dataset. We have also presented a review of approaches for the transformation of compositional data and imputation that can fit datasets of possible interest well. We showed that feature space transformation, denoting differences in the abundances of bacteria, made by all and pivot log ratios, positively affects the accuracy of classification. These results complement the studies of (Karwowska et al., 2025) that used centered and isometric log ratios and concluded that there was no significant improvement in classification performance. We have demonstrated that features showing differences in bacterial abundance or in time are among the keys in prediction tasks. The second key is the imputation technique for missing data.

We found that the imputation techniques yielding the highest classification accuracy for our datasets were Gaussian process regression (GPR) (Williams and Rasmussen, 1995) and conditional generative adversarial networks (cGANs) (Mirza and Osindero, 2014). The choice of the model for imputation affects the learned relationship between incomplete data and time points to fill in. The importance of imputation when some time points are missing can be considered from different perspectives: (i) we increase the size of the training set, including both collected and artificial time points; (ii) we learn on more diverse data, taking into account both complete and incomplete data points; and (iii) we may balance the dataset, including incomplete data points with a particular label, into the analysis. When the dataset is imbalanced, better accuracy is expected to be obtained for a more well-represented dataset. However, in some applications, including classification tasks for two considered datasets, sensitivity may have greater importance than specificity. The imputation and selection of features distributed in time are possible because of the specific format used for this study, where measurements taken at several time points constitute a dataset. For the purpose of imputation, we investigate different models that describe the relationships between time points and use balanced accuracy as a metric for suitability for such models.

We analyzed two datasets. We forecast PPD via gut microbiome data collected before childbirth. We achieved a balanced accuracy of 0.62 and a sensitivity of 0.39, statistically outperforming random guessing and surpassing the accuracy of predictions on the basis of depression states observed during pregnancy. That is, we observed the predictive power of microbiome data for determining PPD. The result is obtained by moving from compositional data to a real space via pivot log ratios (Fišerová and Hron, 2011). All features used for classification are log ratios between species abundances. Even when a time point after childbirth is available, features important for classification are not limited by bacterial abundance measured at the last time point. We increased the value of sensitivity by imputing missing time points with only positive labels. An alternative approach to prioritize sensitivity would involve classification algorithms with guaranteed error control, as explored in (Care, Ramponi, and Campi, 2018).

The results of other articles in predicting postpartum depression (Andersson et al., 2021; Tortajada et al., 2009; Y. Zhang et al., 2021) are higher than our forecasting accuracy. Notably, the last information used therein is obtained at the time of childbirth, while our feature space of interest is limited in time by the 30th week of pregnancy. A fair comparison would require microbiome data collected shortly after delivery. However, the approaches may complement each other. For example, combining microbiome data with additional background and infant-related variables could improve predictive accuracy.

For the prediction task for food allergies in infants, we obtained a balanced accuracy of 0.72, with a sensitivity of 0.66. In original work on the same dataset (Martin, Virkud, Dahan, et al., 2022), the authors obtained a balanced accuracy of 0.66 using time points after the 6th month of infants’ life, while we considered the 6th month as the last time point in the analysis. The abundance collected at all time points is important for the prediction. We obtain the value of accuracy via imputation. Imputation allows the use of incomplete data with real-time points during training.

An increase in accuracy was possible when we limited the input space to the data points where less than half of the data were missing. While we assumed the same models for complete and incomplete data during imputation, further research could explore relaxing this assumption. We worked with datasets where the dimensionality was greater than the size. Thus, we focused on dimensionality reduction. We merged bacteria with high sparsity, selected features and selected the optimal number of features. In such a way, we decrease the feature space to bacteria abundance or their ratios related to labels of interest. Future research could take the inverse approach by limiting the analysis to bacteria with known correlations to the labels and then applying log-ratio transformations and imputation techniques. While the results shown in this paper are both microbiome datasets, the method is applicable to any compositional dataset, including (meta-)transcriptomics and untargeted metabolomics.

We have applied imputation for complementing data, increasing their diversity and size, and for balancing one of the datasets. Future work should focus on oversampling techniques applicable to the considered data structure. By over-sampling, that is, modeling compositional data conditional on labels, researchers can mitigate imbalances in the data, such as rare conditions or underrepresented demographics.

## A Alternatives for the classification algorithms

In the main text, we apply ANOVA to select meaningful classification features. The option of using dimensionality reduction via feature selection is justified here. We first test the (hierarchical) order of features suggested by a hierarchical tree while applying the pivot log ratio. We also consider the order given by Lasso regularization and by principal component analysis (PCA) in Table 9. PCA applied to CLR provides another way to transform compositional data called the isometric log ratio (Egozcue et al., 2003). Tables 10 and 11 present the results when other classifiers are applied, namely, support vector classification (SVC) and adaptive boosting (AdaBoost). The AdaBoost classifier is based on a decision tree, but then it reweighs data points depending on the correctness of the classification.

**Table 9:** Accuracy of feature ordering via the random forest classifier for Dataset 1.

| order | balance | metric | acc | p-value | sens. | spec. | n |
| --- | --- | --- | --- | --- | --- | --- | --- |
| PCA | off | median | 0.54 | 0.01 | 0.18 | 0.9 | 1,2,4,3,2 |
|  | on | mean | 0.55 | 4e-5 | 0.19 | 0.94 | 1,1,18,6,5 |

If the balance is on, the class weights are inversely proportional to the class frequencies. Only results with p-value less than 0.05 are shown. $n$ is the optimal number of features used.

**Table 10:**
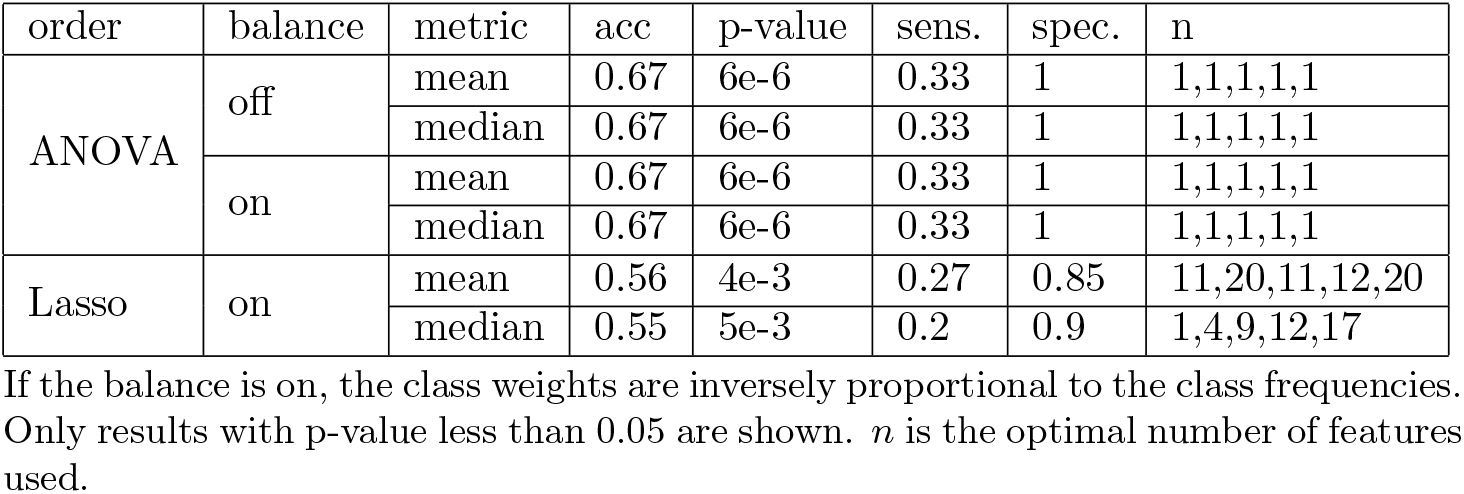
Accuracy for different types of feature ordering using a support vector classifier.

| order | balance | metric | acc | p-value | sens. | spec. | n |
| --- | --- | --- | --- | --- | --- | --- | --- |
| ANOVA | off | mean | 0.67 | 6e-6 | 0.33 | 1 | 1,1,1,1,1 |
|  |  | median | 0.67 | 6e-6 | 0.33 | 1 | 1,1,1,1,1 |
|  | on | mean | 0.67 | 6e-6 | 0.33 | 1 | 1,1,1,1,1 |
|  |  | median | 0.67 | 6e-6 | 0.33 | 1 | 1,1,1,1,1 |
| Lasso | on | mean | 0.56 | 4e-3 | 0.27 | 0.85 | 11,20,11,12,20 |
|  |  | median | 0.55 | 5e-3 | 0.2 | 0.9 | 1,4,9,12,17 |

If the balance is on, the class weights are inversely proportional to the class frequencies. Only results with p-value less than 0.05 are shown. $n$ is the optimal number of features used.

**Table 11:** Accuracy for different types of feature ordering via adaptive boosting.

| order | balance | metric | acc | p-value | sens. | spec. | n |
| --- | --- | --- | --- | --- | --- | --- | --- |
| hierarchical | off | median | 0.54 | 0.01 | 0.2 | 0.89 | 8,20,2,12,7 |
| ANOVA | off | mean | 0.68 | 6e-5 | 0.47 | 0.9 | 5,20,13,5,1 |
|  |  | median | 0.68 | 2e-7 | 0.4 | 0.95 | 6,1,13,2,4 |
| PCA | off | mean | 0.56 | 6e-4 | 0.27 | 0.86 | 1,3,17,16,8 |

Only results with p-value less than 0.05 are shown. The class weights are equal for the chosen classification method. $n$ is the optimal number of features used.

PCA, Lasso, and hierarchical ordering do not improve the accuracy. The results for the hierarchical order are as expected, as the important features are not selected in the same way as in the ANOVA method. As seen in the tables, different classification methods do not lead to an increase in balanced accuracy. From the comparison analysis, we find that adaptive boosting classifiers can improve the sensitivity of the classification when the number of features is constrained.

## Data availability

Dataset 1 is available from NCBI under BioProject PRJEB62678. Dataset 2 can be found on NCBI BioProject PRJNA730851.

## Author Contributions Statement

Conceptualization: A.Sh., C.W., D.Z., L.H., P.S. Data curation: B.T., A.Sk., L.H. Formal analysis: A.Sh. Funding acquisition: C.W., D.Z., L.H., P.S. Investigation: A.Sh. Methodology: A.Sh., C.W., D.Z., L.H., P.S. Software: A.Sh. Resources: A.Sk. Supervision: L.H., P.S. Visualization: A.Sh. Writing – original draft: A.Sh. Writing – review & editing: C.W., D.Z., L.H., P.S.

## Competing interests

No competing interest is declared.

## Acknowledgments

This work was supported by funding from the Dept. of Information Technology, Uppsala University, through equal opportunity research in the IT initiative. P.S. acknowledges support from the Swedish Research Council through grant agreement no. 2023–05593 and the Knut and Alice Wallenberg foundation through the Program for Academic Leaders in Life Science (PALS). This work was partially supported by the SciLifeLab & Wallenberg Data Driven Life Science Program (grant: KAW 2020.0239) and Swedish Research Council contract no. 2024–03903.

## Notes

### Competing Interest Statement

The authors have declared no competing interest.

### Summary of Updates

Authors list is updated together with contribution statements. Several references into introduction are added.

